# Including the Invisible Fraction in Whole Population Studies: a Guide to the Genetic Sampling of Unhatched Bird Eggs

**DOI:** 10.1101/2023.06.13.544703

**Authors:** Fay Morland, Selina Patel, Anna W. Santure, Patricia Brekke, Nicola Hemmings

## Abstract

Early embryo mortality has recently been proven to be a significant component of avian reproductive failure. Due to the difficulty in distinguishing eggs which have suffered early embryo mortality from unfertilised eggs, this cause of reproductive failure has historically been underestimated and overlooked. We describe methods for recognising and collecting undeveloped, unhatched eggs from wild bird populations, identifying and isolating embryonic material in unhatched eggs, and efficiently extracting DNA from those samples. We test these methods on unhatched eggs collected from the field which have undergone post-mortem incubation. We obtained DNA yields from early-stage embryos that are sufficient for a wide range of molecular techniques, including microsatellite genotyping for parentage analysis and sex-typing. The type of tissue sample taken from the egg affected downstream DNA yields and microsatellite amplification rates. Species-specific microsatellite markers had higher amplification success rates than cross-species markers. We make key recommendations for each stage of the sampling and extraction process and suggest improvements potential and protocol modifications. Genetic and possibly genomic analysis of embryos that die early in development has the potential to advance many fields. The methods described here will allow a more in-depth exploration of the previously overlooked causes of early embryo mortality in wild populations of birds, including threatened species.

## Introduction

Long-term, individual based field studies of wild animals are recognised as a valuable asset to research (Taig-Johnston et al., 2017). Observing populations over multiple years and/or with individual level detail, provides unique insight into processes such as demography, aging, selection and climate change, and contributes to various fields, from conservation (Margalida, 2017) to ecology and evolution (Clutton-Brock & Sheldon, 2010). Often, long-term studies involve exhaustive genetic sampling of all individuals to allow pedigree construction, facilitating our understanding of topics such as life-history, quantitative trait variation, adaptation and inbreeding depression (Pemberton, 2008). However, long-term studies often suffer from a phenomenon known as the ‘invisible fraction’ (Grafen, 1988); a subset of individuals that are unaccounted for because they die before they are sampled. Depending on the size of the invisible fraction, their exclusion from studies could have significant implications for our understanding of demographic and evolutionary processes (Grafen, 1988).

In wild bird populations, there is evidence that the invisible fraction may be substantial (Hemmings & Evans, 2020; Marshall et al., 2023; Savage et al., 2022). Birds provide an ideal system for identifying individuals that die early, since each embryo is contained within a hard-shelled egg, outside of the mother’s body, which can be easily examined and sampled. Despite this, early embryo mortality, defined here as the death of an embryo before it is visible to the naked eye, is often unaccounted for in avian population monitoring. This is either because non-developed eggs are assumed to be unfertilised (Birkhead et al., 2008) or because they are not sampled for molecular analysis. For example, studies which use molecular techniques such as sex determination or parentage analysis to study differential embryonic mortality between the sexes (Brekke et al., 2010; DuRant et al., 2016; Eiby et al., 2008; English et al., 2014) or between within-pair and extra-pair offspring (Whittingham & Dunn, 2001) are often missing the invisible fraction of early failed embryos in their assessment of primary sex ratio.

The inclusion of early-dead embryos in genetic analyses of bird populations is hampered by technical challenges, the most significant of these being DNA degradation. In most long-term studies or monitoring programmes, unhatched eggs remain in the nest until the end of the incubation period, to reduce the risk of mistakenly removing a viable egg or the intervention leading to nest abandonment. This is particularly important for species of conservation concern. However, the microclimate inside a bird’s nest during incubation provides the ideal environment for post-mortem DNA degradation. Although nest temperatures during incubation vary depending on species, habitat and nest characteristics (e.g. 15-26°C in northern flickers (Wiebe, 2001), 35°C in ostrich (Swart et al., 1987), 28-52°C in horned lark (Hartman & Oring, 2003)), nests are generally kept warm by incubating parents to enable embryonic development. This warm environment is likely to accelerate DNA degradation in deceased embryos by facilitating autolysis (Williams et al., 2015). Despite this, DNA samples taken from deceased late-stage embryos, which have been incubated post-mortem, have been successfully used for genetic sex determination (Cichoń et al., 2005). Genetic analysis of early embryos has also been demonstrated using samples taken from live embryos at 1-4 days of development (Strausberger and Ashley, 2001) and embryos that died at an early stage of development and were then incubated for up to 2 days post-death (Kato et al., 2017). However, the successful extraction and molecular analysis of DNA from embryos in unhatched eggs that have died early and undergone a significant period of post-mortem incubation, as is often required by restricted data sampling regimes of wild/protected species, has not been reported. Methods outlining tissue sampling for DNA extraction of early-stage embryos that have undergone full-term post-mortem incubation will prove useful for researchers hoping to sample unhatched eggs from wild bird populations, since the ability to use such samples provides access to a demographic group that has thus far been missing from genetic and demographic studies of wild populations.

Here we describe field methods for collecting and storing fully incubated, undeveloped eggs for subsequent dissection. We also describe methods for obtaining cell samples from embryos that died very early in unhatched eggs, and for extracting DNA from those samples. We apply our methods to fully incubated unhatched eggs sampled from a wild population of birds and assess the quantity of DNA obtained and its suitability for microsatellite genotyping, sex determination and paternity analysis, compared to blood samples taken from live individuals from the same species.

### Protocol Recommendations and Special Considerations for Early Embryo Samples

#### i) Sample Collection and Storage

When collecting unhatched eggs for analysis from wild nests, it is vital to avoid jeopardising the success of the breeding attempt. This is of particular importance when working with species of conservation concern. We recommend candling eggs no earlier than 4 days after the onset of incubation to identify non-developing eggs (i.e., those with no sign of embryo development) before removal. The technique of candling involves shining a bright light through the egg by placing a torch adjacent to a held egg or an egg within the nest to look for signs of development, i.e., dark spots and blood vessels. In protected species, undeveloped eggs may need to be left in the nest until after any other eggs have hatched (or a few days beyond the incubation period, if no eggs hatch). In either instance, to limit DNA degradation after collection, we recommend unhatched eggs are stored at 1-5 °C as soon as possible. Samples can be placed in a cool box when in the field and afterwards transferred to a standard refrigerator where they should remain until processing.

#### ii) Sample Processing and Cell Isolation

Unhatched, undeveloped eggs are usually found in two distinct states when collected from the field: either the yolk remains intact and separate from the albumen inside the shell (referred to here as separated; Fig. 1.a), or the contents are addled (yolk broken down and mixed with albumin, often degraded to some degree (Fig. 1.c)). This distinction has been used in the past to falsely categorise eggs as unfertilised when they are separated, or as failed embryos when they are addled (e.g. Cook et al., 2005). However, separated eggs may be fertilised and contain embryonic cells (Assersohn et al., 2021; Birkhead et al., 2008).

**Fig. 1.**
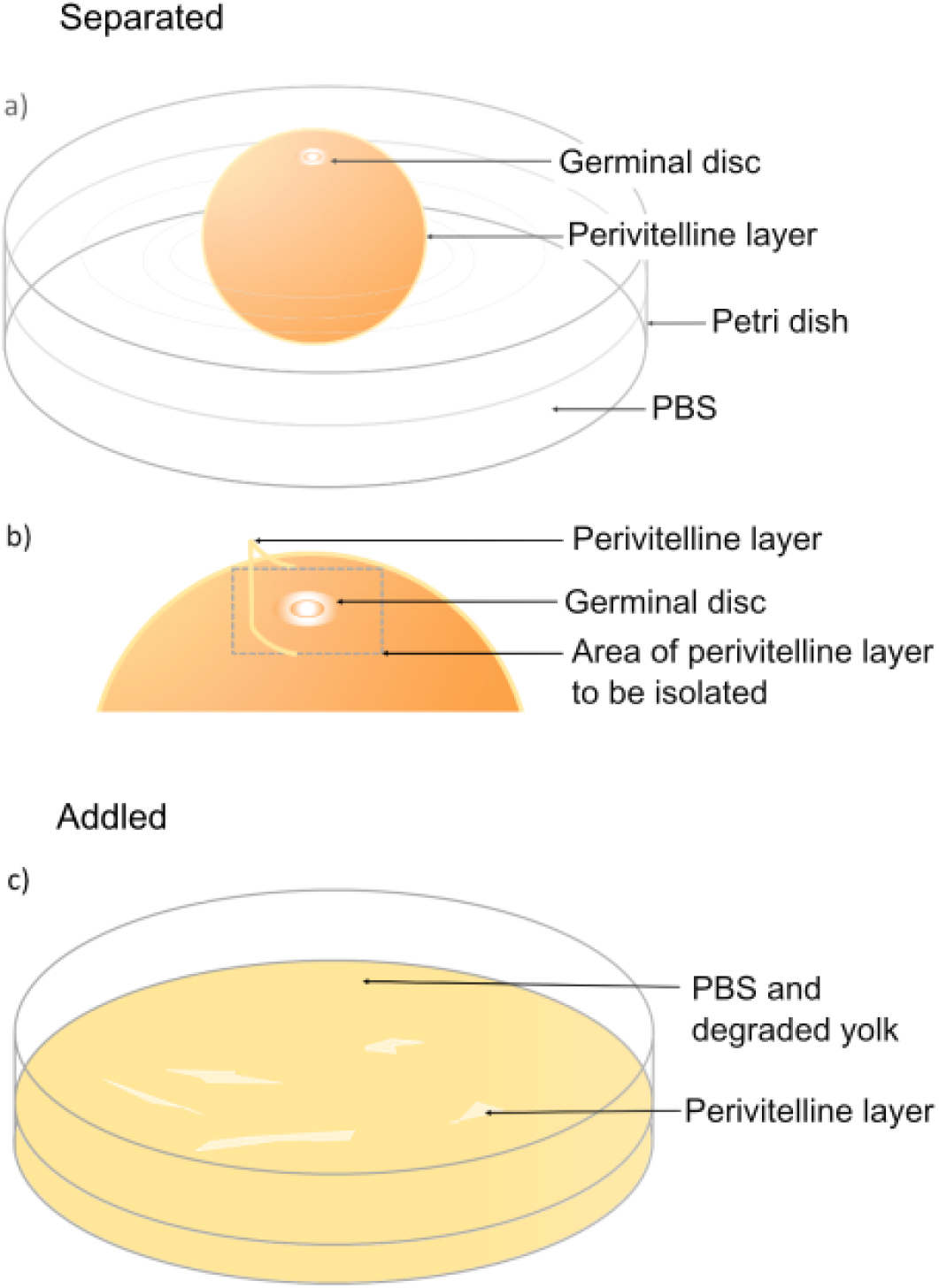
a) A separated unhatched egg removed from the shell and placed in phosphate buffered saline solution (PBS) for dissection and isolation of two key parts: the germinal disc and the perivitelline layer. b) Isolating the germinal disc and the area of perivitelline layer surrounding the germinal disc will provide the best opportunity of locating embryonic cells and sperm cells. c) An addled unhatched egg removed from the shell and placed in PBS will create a cloudy mix of yolk, PBS and pieces of degraded perivitelline layer. Locating and isolating pieces of perivitelline layer, on which embryonic cells may be attached and in which sperm cells may be trapped, provides the best opportunity of finding evidence of fertilisation and embryonic development in addled unhatched eggs.

For a detailed explanation of the equipment and methodology used to assess the fertility status of unhatched eggs, please see Assersohn et al. (2021) and the open access protocols and videos referred to therein (“Practical resources for identifying the causes of hatching failure in birds” 2021). In brief, to assess an undeveloped egg for signs of fertilisation and embryonic cells, first open the egg into PBS (phosphate buffered saline, pH 7.4) in a petri dish (Fig. 1). Secondly, locate the parts of the egg most likely to contain embryonic cells and sperm cells: the germinal disc and the section of perivitelline layer surrounding it. The germinal disc, which contains the female pronucleus, is the site of fertilisation and where embryonic development starts. It appears as a white spot or ring on the surface of the yolk of intact eggs (Fig. 1.a and Fig. 1.b). The perivitelline layer consists initially of a single layer (the ‘inner’ perivitelline layer), which surrounds the yolk and is penetrated by sperm prior to fertilisation. Shortly after ovulation, a second perivitelline layer (the ‘outer’ layer) is formed through the secretion of glycoproteins from the oviduct, and sperm that are around the egg at that time are trapped in this outer layer as it forms. Sperm cells can therefore be found in the perivitelline layer, and embryonic cells can be found either in the germinal disc or attached to the overlying perivitelline layer. By locating, isolating, and examining the germinal disc and the perivitelline layer, following the methods of Assersohn et al. (2021), it is possible to determine whether sperm reached the egg, and whether fertilisation and embryonic development took place.

When the egg is separated (Fig. 1.a), locating and isolating the germinal disc macroscopically is straightforward. However, when the contents of failed eggs collected from the field are heavily degraded or “addled” (Fig. 1.c), searching the sample macroscopically or using a stereo microscope to find pieces of perivitelline layer provides the best opportunity of locating the germinal disc and embryonic cells. For separated eggs: once the germinal disc has been located, cut and remove the surrounding section of perivitelline layer, using surgical scissors and tweezers, and place it into a clean dish of PBS. Remove the germinal disc from the yolk using a pipette or a hair loop (a loop of human hair taped to a pipette tip which creates a delicate tool) and place straight onto a microscope slide. For addled eggs, remove all pieces of perivitelline layer into a clean dish of PBS. For pieces of perivitelline layer isolated from both intact and addled eggs, clean off any excess yolk with a hair loop or by agitating in PBS solution. However, take care not to clean so thoroughly as to potentially dislodge any embryonic cells which may be attached, i.e., remove large pieces of yellow yolk which may obscure the perivitelline layer under the microscope, but do not remove all material on the perivitelline layer, as some of it may be embryonic cells. Place the perivitelline layer onto a microscope slide using tweezers and flatten/smooth the pieces out using the hair loop. Once the germinal disc and perivitelline layer are on the microscope slide, they should be stained with the fluorescent DNA dye Hoechst 33342 and examined with a fluorescence microscope to search for embryonic cells, which will appear as clustered fluorescent-blue cells (Fig. 2). If embryonic cells have been located on the germinal disc or the perivitelline layer, transfer the whole sample from the microscope slide into absolute ethanol to be stored for later DNA extraction.

**Fig. 2.**
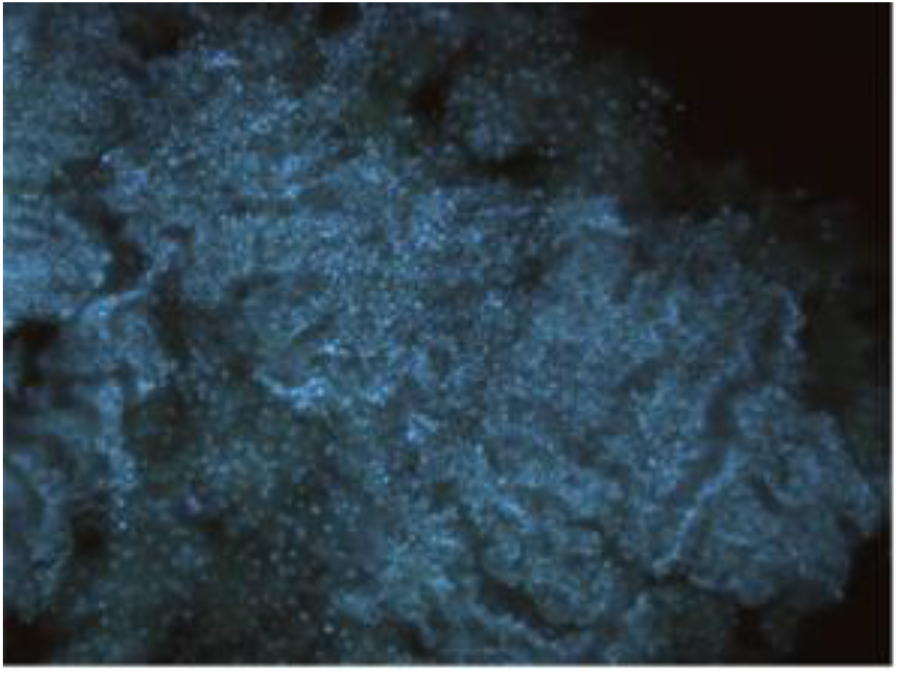
Hihi (*Notiomystis cincta*) embryonic cells stained with Hoechst fluorescent dye, as viewed at a magnification of 100x (10x objective lens X 10x eyepiece lens). Embryonic cells are shown as bright blue clusters of cells against a background of unstained yolk/PBS, appearing black

**Fig. 3.**
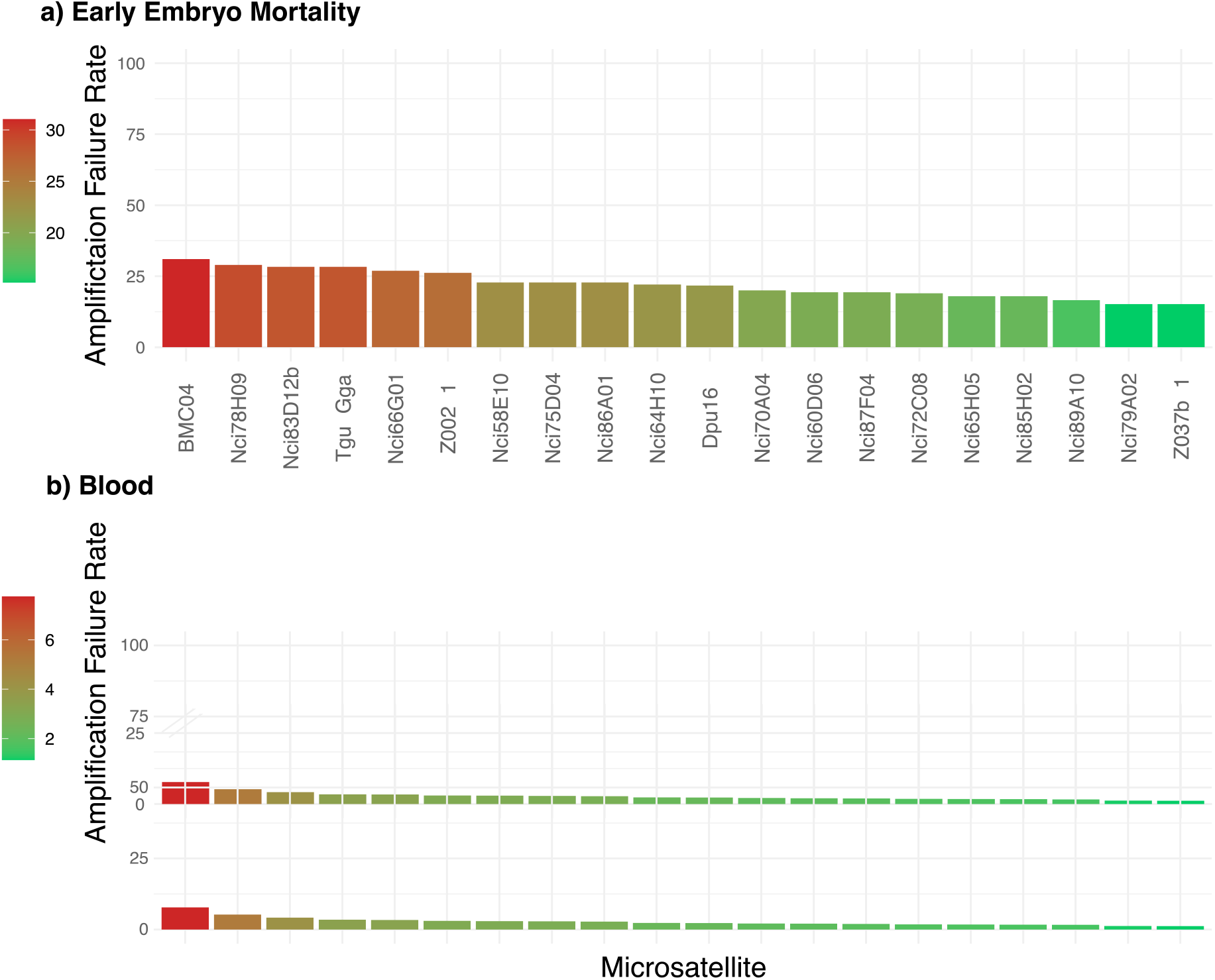
The microsatellite amplification failure rates for DNA obtained from a) cell samples taken from embryos that died early (n=150) and b) blood samples taken from nestlings (n=3,162). The green to red colour gradient indicates the markers which failed the least (green) to markers which failed the most (red), relative to the failure rates for each sample type; note the different scales of the colour gradients. Cross-species markers are indicated by *. The two sex-linked markers are Z002 and Z037b

#### iii) DNA Extraction

Some samples from embryos that died at an early stage of development contain a very small number of cells. It is therefore necessary to take specific steps to maximise the amount of tissue retained for the DNA extraction. We found that the following modifications to the Qiagen DNeasy Blood and Tissue kit spin-column protocol for tissue samples (Purification of Total DNA from Animal Tissues (Spin-Column Protocol)) improved DNA yield. Instead of starting step 1 of the protocol by placing tissue in a separate microcentrifuge tube, the ethanol-preserved perivitelline layer and germinal disc samples should be centrifuged in their original sample tubes at maximum speed (13,000 RPM) for 1 minute. All ethanol should be removed using a pipette, taking care not to disturb the tissue pellet at the bottom. After ethanol removal, 180ul Buffer ATL and 40ul Proteinase K should be added directly to the sample tubes. Sample tubes should then be incubated on a rocking platform at 56°C for one hour. Only after this step should the samples be transferred into new 1.5 ml Eppendorf tubes, followed by incubation in a rotating oven at 56°C overnight or until the samples are completely lysed. The method of lysis in the original sample tube was adopted to ensure that no cells (e.g., those potentially attached to the walls of the sample tube), were lost through transferring the sample into a new tube for extraction. Following this modification, the Qiagen Blood and Tissue kit protocol can be followed without further modifications from steps 3 to 7. Following extraction, the concentration of the DNA should be tested and samples concentrated/diluted to a standard concentration required for amplification and genotyping. While we used the Qiagen DNeasy Blood and Tissue kit, other extraction kits or protocols are likely to also prove successful with similar modifications to reduce sample loss.

## Methods: Case study

### i) Study system

The eggs we used to test our methods were collected from a managed population of hihi, *Notiomystis cincta*, which nest in nest boxes on Tiritiri Mātangi Island, Hauraki Gulf, Aotearoa (New Zealand). This population is monitored during the breeding season, and fledglings are colour banded and blood sampled, allowing for individual identification and the construction of a long-term genetic pedigree using microsatellite genotyping (Brekke et al., 2009, 2015). Due to the high rates of extra-pair paternity in this species, genotyping is required for accurate parentage assignment (Brekke et al., 2013; Ewen et al., 2004). For the purposes of this study, unhatched eggs with no sign of embryonic development were collected from this population across two breeding seasons spanning 2019-2021, screened for signs of embryonic development, and sampled for DNA. The hihi is an IUCN listed vulnerable species and exists only in predator-free reserves in the North Island of Aotearoa (Brekke et al., 2011). As is typical for many protected bird species, there are limitations to the amount of nest interference that is permitted during the hihi incubation period. Hihi have a 14-day incubation period, and unhatched eggs were collected on day 13, to limit disturbance to natural incubation behaviour and comply with permits. Before collection, eggs were candled at the nest box with a long torch to ensure there was no embryonic development visible. A long torch was used because it allows eggs to be candled in situ, avoiding the need to handle or remove eggs from the nest. In the field, undeveloped eggs were stored in small, sealable sample bags and placed inside plastic 50mL centrifuge tubes. The sample bags fitted closely into the tube and buffered the egg from movement and impact during transportation to the field station/lab. The centrifuge tubes containing eggs were placed into a cool box with ice blocks for transportation from the field site to a fridge to minimise further tissue and DNA degradation as much as possible. The samples were then examined within 19 days of collection using the microscopic methods described in the Protocol Recommendations (section ii). If embryonic development was visible macroscopically or via a stereomicroscope, the embryo’s developmental stage was classified using the Hamburger-Hamilton (HH) embryonic development staging series (Hamburger & Hamilton, 1951) and a tissue sample was taken. 145 tissue samples were taken from eggs found to contain embryonic cells, 49 samples were from the first season (2019-20) and 96 were from the second season (2020-21). The tissue samples were stored in ethanol before DNA extraction using the methods described in the Protocol Recommendations (section iii). The blood samples used as a comparison were taken before fledging, from the 21 mothers of the embryos included in this study, between the years of 2013 and 2018.

### ii) Microsatellite Analysis

DNA was quantified using a Qubit fluorometer and diluted if over 25ng/μl or concentrated if under 5ng/μl using a DNA vacuum concentrator, depending on the initial concentration, and volume of the DNA. The target concentration for this microsatellite analysis method when using blood samples is 20ng/μl, however due to the low concentration and volume of some samples, and to avoid excessive drying and re-diluting which can cause DNA damage and salt accumulation which can inhibit PCR, we aimed for a sample concentration between 5 and 25ng/μl. The average initial concentration was 20.5ng/μl. DNA samples from embryos that died early in development and remained in the nest for the full incubation period were genotyped using 20 microsatellite markers, comprising of 2 sex-typing markers (Z002a and Z037b; Dawson, 2007; Dawson et al., 2015), 15 species-specific markers, and 3 markers developed for other passerines (Brekke et al., 2009, two primers listed in this paper were not used: Nci014 and MSLP4).

Microsatellite regions were amplified using PCR before genotyping, with each 6.75μL PCR containing 2μl of genomic DNA (at the average concentration 20.5ng/μl this resulted in each PCR reaction containing an average of 41ng of DNA), 3μl of Qiagen PCR Kit Mix and 1.75μl of 1 of 4 multiplex primer mixes. Each multiplex primer mix contained 4-6 primers at 0.2 - 0.6μM concentration. Primers were grouped into multiplex mixes based on annealing temperature and allele size range (Brekke et al., 2009). The PCR followed a thermal cycle of 9°C for 15 minutes, followed by 30 cycles of 94°C for 30 seconds, annealing temperature of 56°C or 64°C (depending on primer mix, see Brekke et al., 2009) for 30 seconds, 72°C for 90 seconds, and a final step of 72 °C for 10 minutes. Microsatellites were genotyped using an Applied Biosystems 3130 Genetic Analyser and output used to assign microsatellite genotypes for each individual and locus using GeneMapper 5 software. The same microsatellite analysis protocol is used for genotyping fledglings using DNA obtained from blood samples, although Geneious software with the Microsatellite plugin is sometimes used for genotyping. Due to COVID-19 travel restrictions, the genotyping of the 2020-21 season was performed in a different laboratory and on a different sequencer to the other samples included in this study and to the laboratory and sequencer that was used to characterise the population’s alleles.

### iii) Parentage Analysis

To test the ability of the resultant microsatellite genotypes to accurately infer known maternal links, the sample genotypes were used to carry out parentage analysis in Colony software (Jones & Wang, 2010). Colony is provided with the genotypes of the offspring (here, egg samples as well as other fledglings from the same sampling year) and of all candidate mothers and fathers. In addition, information can be provided about locus error rates. Information can also be provided about the paternal and/or maternal sibship of the offspring and known mothers and fathers. In the case of hihi, paternal sibship and paternity cannot be inferred due to the high rates of extra pair paternity (Brekke et al., 2013). However, maternal sibship (individuals from the same nest) and maternity can be accurately inferred from field data due to the individual identification of nesting individuals from their unique colour band combinations.

In an initial Colony analysis with maternal sibship and known maternal ID included, it became clear that there were high rates of non-concordance between the failed embryo and the maternal genotype i.e., the offspring genotype shared no alleles with the maternal genotype at a given locus, indicative of mistyping or allelic drop-out. Further investigation revealed that this was likely a result of mis-calibration of allele sizes across different sequencing laboratories. Sequencing outputs were manually rechecked with peak size assignment adjusted and genotypes re-called before estimating locus specific error rates (due to e.g. mutation or allele dropout) based on persistent non-concordance with known maternal genotypes. An error rate of 0.01 (1%) for all microsatellite markers is used when performing parentage analysis on fledgling blood samples (Brekke et al. 2015), however we expected higher error rates here given that these samples are likely to be degraded. A study on the effectiveness of including markers with high error rates in paternity analysis found that markers with error rates up to 70% are still informative, provided the correct error rates are provided to Colony (Wang, 2019).

Colony was then run with the following parameters: a “full-likelihood” analysis method, “high” likelihood precision, “weak” sibship priors and with locus specific genotyping error rates (range 3%-16%). We ran Colony both with (i) maternal sibship included but no maternal ID and (ii) with maternal sibship and maternal ID included, to (i) simulate the application of this method to field sampling where all eggs or offspring in a nest are sampled but parents may not be sampled or known and (ii) to utilise the known mother to estimate mother-offspring genotype concordance, and hence estimate genotyping error in the degraded egg sample. Colony outputs include the reconstructed pedigree and the probabilities associated with each of the reconstructed parent-offspring links. Before running Colony, we also checked for genotype duplicates between an egg sample and a parent, which may indicate parental DNA contamination of the egg sample.

## Results

### i) DNA Quantity

Most of the 145 undeveloped egg samples (142) that were found to contain microscopically visible embryonic cells, yielded enough DNA following extraction using the methods described in the Protocol Recommendations (section iii) for detection using a high sensitivity Qubit assay. 3 samples had DNA concentrations too low to be read (and are included in mean estimates as a concentration of 0), however upon genotyping only 2 of these samples failed to amplify at any microsatellite, suggesting that only 1 of the 145 samples yielded no DNA. The mean total DNA quantity obtained per egg from the samples was 2,620ng (n=145), but values for individual samples ranged from 6.6ng to 12,400ng. For comparison, the expected yield of DNA from nucleated bird blood when using the Qiagen DNeasy Blood and Tissue kit is 9,000 – 40,000ng (“DNeasy Blood & Tissue Kits,” Accessed: 19-09-2022). The yield of DNA obtained varied between embryo stages and sample type (Table 1). As expected, the quantity of DNA obtained was higher for embryos that died at a later stage of development, i.e., those that didn’t reveal development on candling but an embryo was obvious via macroscopic inspection once the egg was opened. For very early embryonic samples ≤HH stage 7), where no development was visible on macroscopic examination, samples of perivitelline layer with the germinal disc attached provided the highest yields of DNA on average. The results of an analysis of variance including 93 very early embryonic embryo samples (≤ HH stage 7) with both sample type and DNA yield recorded show that sample type significantly impacts the DNA yield (F = 6.3, p = <0.05, df = 2), and a post hoc Tukey test revealed that samples of the germinal disc yield significantly more DNA than samples of perivitelline layer (diff = 2.926ng, p = <0.01).

Samples were either kept in the refrigerator before microscopic analysis and DNA sampling for between 2 and 19 days, or frozen on the day of removal from the nest and frozen for between 361 and 463 days before microscopic analysis and DNA sampling. For refrigerated samples we did not find a relationship between processing latency (i.e., the time between egg collection and dissection/storage of sample in ethanol) and DNA quantity (cor = 0.06, p = 0.7, df = 40) or the number of microsatellites amplified in downstream genotyping (cor = 0.012, p = 0.9, df = 42).

We did not find a significant difference in DNA quantity (F = 0.008, p = 0.93, df = 1), or the number of microsatellites amplified in downstream genotyping (F = 1.67, p = 0.2, df = 1) between samples which were frozen or refrigerated before sampling.

### ii) Microsatellite Analysis

Of all samples processed, 50 out of 145 samples amplified across the entire microsatellite panel and the majority (135 samples) showed at least some amplification. Mean microsatellite amplification rates were similar for pre- and post-HH stage 8 embryos (Table 1) and very poor amplification rates were found only for perivitelline layer samples categorised as having very few embryonic cells found during microscopic examination (average success rate = 10%, n = 6). DNA extracted from the germinal disc/embryonic material provided the provided the highest yields of DNA and the highest success rate in microsatellite analysis. These results suggest that prioritising samples in which the embryo or germinal disc can be found, and discounting samples for which very few embryonic cells are found during microscopic examination, will prove the most efficient approach if resources and/or time are limited.

The average microsatellite success rate was 77% across samples, representing a higher failure rate in microsatellite amplification than blood samples taken from fledglings from the same population, for which the average microsatellite amplification success rate 97.4% (Fig. 2). The DNA amplification showed some of the highest rates of failure with cross-species microsatellite primers; two of the four highest failure rate of the microsatellite markers were two of the three cross-species markers; BMC04 (37.5%) and Tgu-Gga (35.8%). This suggests that genotyping efforts for early embryonic samples may be more productive when using species-specific molecular markers. The length of the target allele was not associated with microsatellite failure rate when including (cor = -0.004, p = 0.9, df = 18), or excluding (cor = -0.18, p = 0.5, df = 15) cross-species primers (Fig. 4b).

**Fig. 4.**
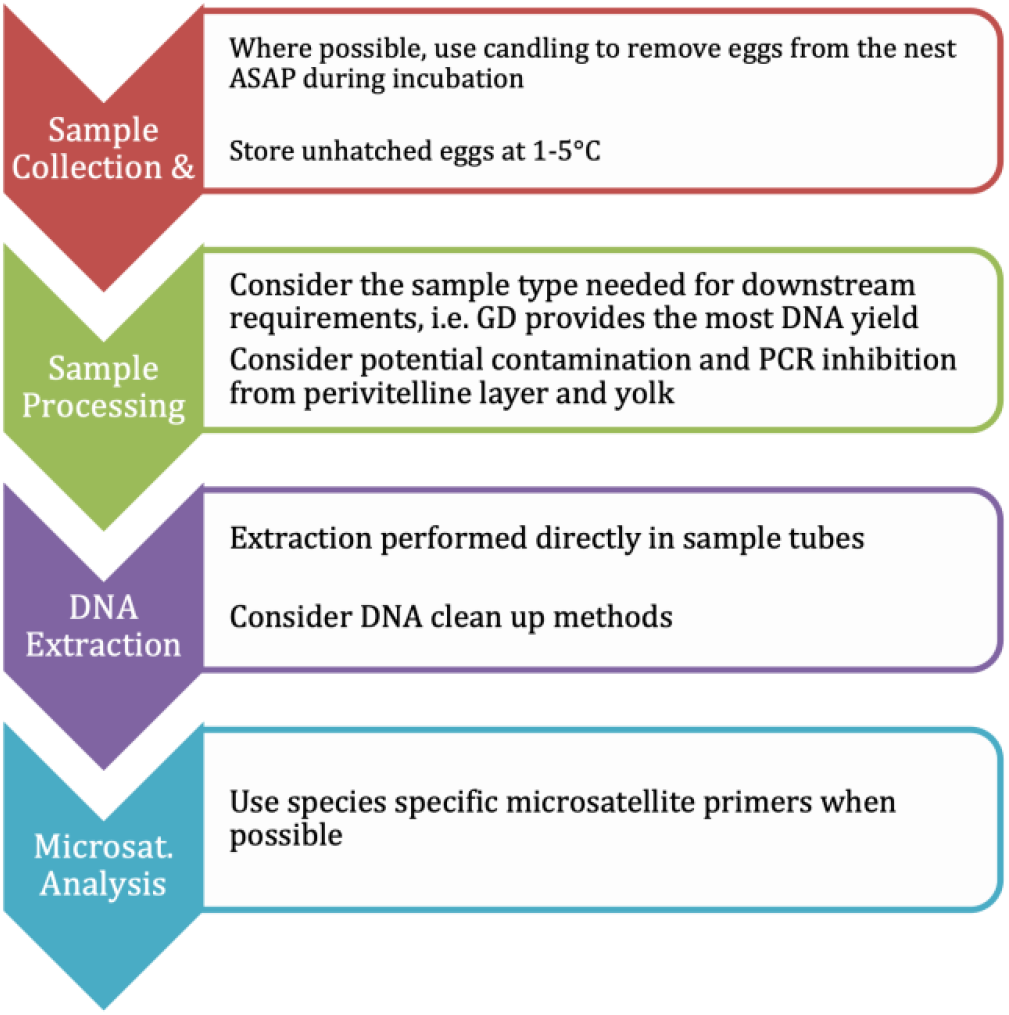
A summary of the important steps recommended during sample collection, storage, processing, DNA extraction and microsatellite analysis.

**Table 2.**
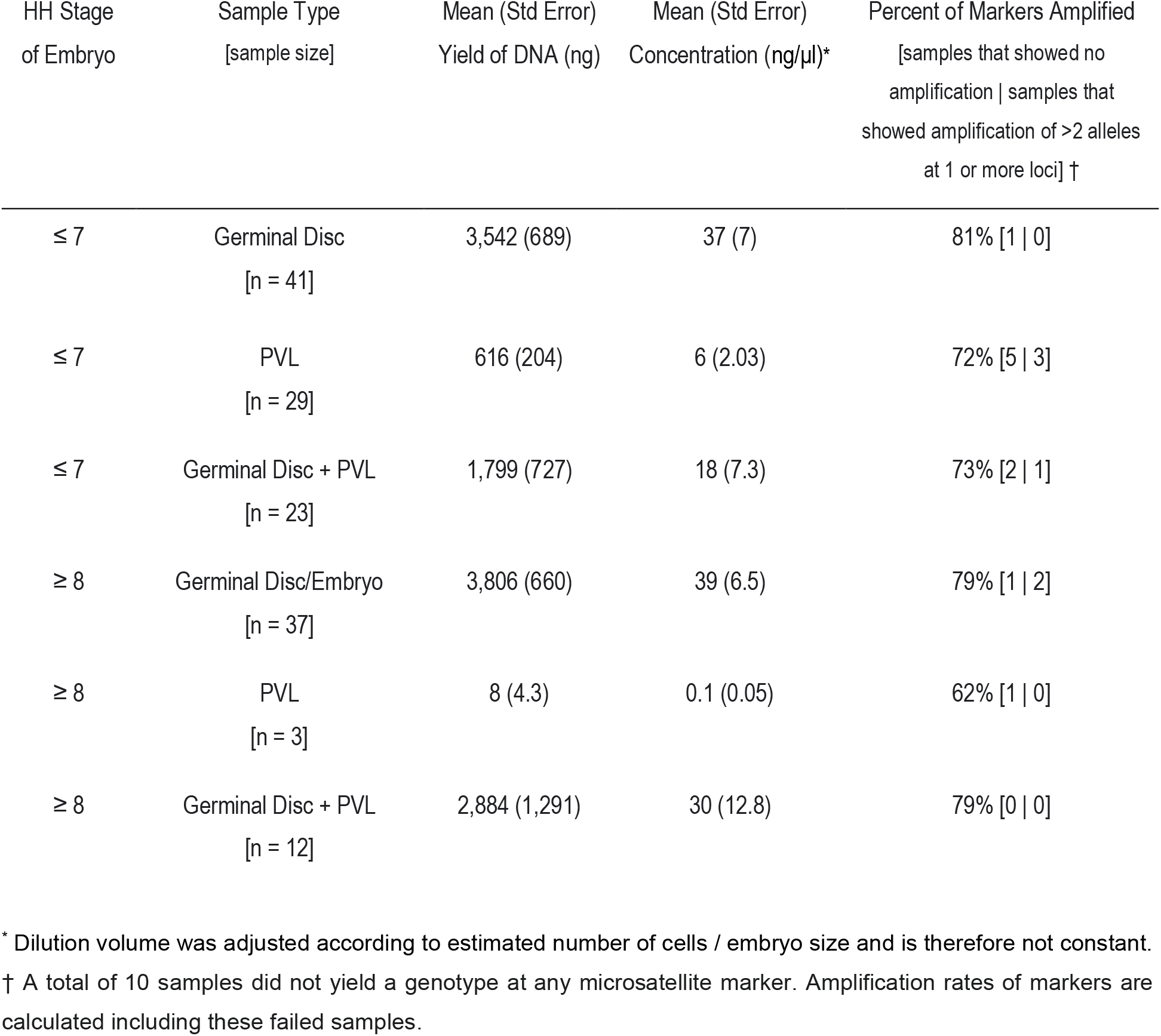
The mean yield of DNA and microsatellite amplification rates from germinal disc, perivitelline layer (PVL), and embryos of different developmental stages (HH Stage refers to Hamburger-Hamilton developmental stages (Hamburger and Hamilton, 1951)).

### iii) Parentage Analysis & Sex Typing

There were 6 potential cases of DNA contamination from parental DNA, detected as individuals amplifying for 3 or more alleles at a single locus. The contamination may have originated from sperm trapped in the perivitelline layer (Carter et al., 2000), maternal and/or paternal DNA from other egg components such as egg shell fragments (Martín-Gálvez et al., 2011; Strausberger & Ashley, 2001) or cross-contamination during sampling or extraction. The 6 samples with potential contamination were not included in Parentage analysis.

Despite attempting to resolve calibration errors between sequencing machines before running Colony analyses, there were also still cases of non-concordance between the offspring and the mothers, with 8.46% of alleles non-concordant with maternal genotypes. Of these non-concordant alleles, 55% were homozygous, suggestive of allelic drop-out, however the cause of the remaining 45% is unclear. Non homozygous alleles which are non-concordant with maternal genotyping could result from technical issues such as laboratory temperature differences (Davison & Chiba, 2003) or electrophoresis artefacts (Fernando et al., 2001), a lack of DNA quality and/or quantity (Goossens et al., 1998) or human error at any stage of the process (Bonin et al., 2004), including human error in field data recording, microsatellite peak assignment or PCR and/or sequencing protocol. In this case, it may also be possible that not all calibration errors between sequencing machines were fully resolved.

Despite the variable genotyping rates across loci and across samples, and the presence of non-concordant genotypes between samples and their known mothers, Colony successfully inferred maternity in 69% of cases where maternal sibship, but not maternal identity, was given and in 74% of cases where both maternal sibship and maternal identity were given.

Most samples (84%) could be sexed from the 2 sex-typing markers used in microsatellite analysis. The sex ratio of all failed early embryos was 1.5 (males:females), which may be due to sex-biased mortality at this stage, or allelic drop-out during microsatellite genotyping leading to incorrect sex assignment. However, the latter explanation is unlikely since although the sex bias was largest in the early stages of embryonic failure (≤ HH stage 7, 1.96 males:females, n=77) compared to the later stages of embryonic failure (≥ HH stage 8, 1.09 males:females, n=48), the rate of identified mistypes was not higher in these early stage samples (≤ HH stage 7, mean number of mistyped alleles = 9) compared to the later stages (≥ HH stage 8, mean number of mistyped alleles = 10). In addition, the use of two separate sex markers may help to protect against sex-typing errors due to allelic drop-out.

## Discussion

The results of our study demonstrate that it is possible to harvest tissue from avian embryos that die early in development and use this for DNA extraction and downstream molecular analyses, even when unhatched eggs are incubated for prolonged periods in natural nests prior to sampling. Our findings suggest this approach may be highly applicable to a wide range of wild bird population studies across multiple fields. With particular care given to sample collection, microscopic fertility examination techniques, and DNA extraction protocols, it is possible to obtain high yields of genomic DNA, even with very early embryos (≤ HH stage 7/less than 2 days development). The DNA we extracted from early embryo samples was of relatively high molecular yield despite small quantities of often degraded tissue available, and it was also found to be suitable for several molecular techniques including microsatellite genotyping for sex typing and parentage assignment.

Fig. 4 highlights the steps that are important for the sampling, processing, DNA extraction and microsatellite sequencing of failed embryos in undeveloped unhatched eggs.

In our study, all but one sample extracted in this study yielded at least some DNA detectable through a high sensitivity Qubit assay. This is a higher success rate than a previous attempt to extract DNA from bird faeces, which had extraction success rates of 34-80% depending on the method used (Alda et al., 2007). In addition, the mean amount of DNA obtained from early-dead embryos in our study was 2,122ng, which is ample for several molecular techniques. For example, a minimum of 10ng of DNA per is recommended per reaction for multiplex microsatellite analysis (Brekke et al., 2010; Narina et al., 2011; Neff et al., 2000), 2.5ng of DNA per sample is required for RAD-seq (Etter et al., 2011), 20-1000ng for llumina Sequencing (“Sample Requirements,” Accessed: 05-09-2022), depending on the precise method, and a maximum of 200ng is required for Sanger sequencing (“Sample Requirements,” Accessed: 05-09-2022).

An advantage of microsatellite analysis is that it can be applied to degraded DNA due to the short sequence length of microsatellite target regions (Queller et al., 1993). We demonstrate here that DNA samples taken from embryos that died early in development and were then left in the nest for up to two weeks had an average microsatellite amplification success rate of 77. Although this is substantially lower than the average amplification success rate of blood samples stored in ethanol immediately after sampling (97% - Fig. 2), it does at least allow some degree of genotyping and downstream analysis, for example parentage analysis and sex-typing. It has been demonstrated that short microsatellite markers are more effective than longer microsatellite markers for degraded DNA, such as that from museum specimens (Nakahama & Isagi, 2017). Our results here suggest that limited DNA degradation may not be an issue in early-dead embryo samples, as we see no relationship between the length of the microsatellite marker and the amplification failure rate. Microsatellite amplification failures may instead be due to the presence of proteins and fats (Acharya et al., 2017), which are known PCR inhibitors (Schrader et al., 2012) and make up a large component of the egg yolk (Kowalska et al., 2021). Another category of known PCR inhibitors are proteases (Schrader et al., 2012), which are also present in egg yolk (Shbailat et al., 2016). A total of 412 proteins, including glycoproteins, have also been identified in the chicken perivitelline layer (Brégeon et al., 2022), which may also act as inhibitors. Samples containing perivitelline layer had the worst rates of microsatellite amplification in this study, which may be a consequence of the large number of proteins present in those samples. An improvement on the methods presented here could involve a DNA purification step, such as ethanol precipitation, silica columns or magnetic bead cleaning, to remove protein and lipid contaminants.

Our results also showed that cross-species microsatellite primers were more likely to fail on early embryo DNA samples compared to species-specific primers. Microsatellite sequencing with cross-species primers has previously been used successfully on DNA obtained from faecal samples (e.g. Wultsch et al., 2014) and the benefits of using cross-species include a reduced time commitment and cost compared to developing and optimising species-specific primers as well as utility across many species (Dawson et al., 2010). However, it is reasonable to expect them to show higher rates of failure due to their lack of complete specificity.

Until recently, early embryo mortality has been largely ignored in avian population studies, and individuals that die early in development have for the most part been omitted from molecular studies. As such, our understanding of the evolutionary processes driving population dynamics has been limited. For example, the only previous study (to our knowledge) that has included early embryos in an assessment of primary sex ratio found that a female-biased secondary sex ratio in Eurasian tree sparrow (*Passer montanus*) was due to differential mortality in very early embryos (≤ HH stage 5), with 97% of early mortalities being males (Kato et al., 2017). Here we have shown that molecular information can be relatively easily gleaned from the “invisible fraction” (Grafen, 1988) of bird embryos that die early, and this information is likely to have important implications for ecology, evolution, and conservation. Specifically, data on the invisible fraction will provide vital information on levels of selection in wild populations, as well as improving our understanding of life-history trade-offs (e.g., sex allocation), reproductive biology (e.g., levels of extra-pair paternity) and inbreeding depression. Genetic and potentially genomic analysis of embryos that die early in development has the potential to inform these and many other areas of study. We hope that the methods described in this paper will be widely adopted, allowing a more in-depth exploration of the nature of the invisible fraction and the causes of early embryo mortality.

## Acknowledgements

We acknowledge Ngāti Manuhiri as Kaitiaki of hihi and the significance of Tiritiri Matangi to iwi whose rohe encompasses the Hauraki Gulf | Tīkapa Moana. We would like to thank hihi conservation officer Mhairi McCready and field assistants Leani Oosthuizen and Emma Gray for data collection support as well as volunteers, past students and Department of Conservation staff who have contributed to monitoring the Tiritiri Matangi hihi population and to the Hihi Recovery Group, Department of Conservation, and Supporters of Tiritiri Matangi for maintaining such a long-term vision in monitoring and management of this population. Permissions to conduct the research and collect failed hihi eggs were granted by the New Zealand Department of Conservation permit numbers 36186-FAU, AK/13939/RES, 53614-FAU and 66751-FAU. We thank Cristina Ariani, Gemma Clucas, Johanna Nielsen, Kang-Wok Kim and Shuqi Wang for support with microsatellite genotyping. Many thanks to Dada Gottelli and Kevin Hopkins of the Institute of Zoology and Adrian Turner of The University of Auckland for support with microscopy and genotyping which formed the basis of this manuscript. This research was financially supported by a Dorothy Hodgkin Fellowship (Royal Society; DH160200) awarded to Nicola Hemmings, Fay Morland’s PhD studentship, provided by a Royal Society Research Grant (RGF\R1\180101) and by Research England and British Birds Charitable Trust Grant to Patricia Brekke and The Royal Society of New Zealand Marsden Fund to Anna Santure. The authors have no competing interests to disclose.

## Author Contributions

FM wrote the manuscript with support from all other authors. NH conceptualised the study. Sample collection, microscopy and data analysis was carried out by FM. NH, PB, FM and SP developed the methods used in the study with support from AS. FM and SP carried out the DNA extraction and sequencing with support and supervision from NH, PB and AS. All authors read and approved the final manuscript.

